# Multimodal 3D Image Registration for Mapping Brain Disorders

**DOI:** 10.1101/2024.08.24.609508

**Authors:** Hassan Mahmood, Syed Mohammed Shamsul Islam, Asim Iqbal

## Abstract

We introduce an AI-driven approach for robust 3D brain image registration, addressing challenges posed by diverse hardware scanners and imaging sites. Our model trained using an SSIM-driven loss function, prioritizes structural coherence over voxel-wise intensity matching, making it uniquely robust to inter-scanner and intra-modality variations. This innovative end-to-end framework combines global alignment and non-rigid registration modules, specifically designed to handle structural, intensity, and domain variances in 3D brain imaging data. Our approach outperforms the baseline model in handling these shifts, achieving results that align closely with clinical ground-truth measurements. We demonstrate its efficacy on 3D brain data from healthy individuals and dementia patients, with particular success in quantifying brain atrophy, a key biomarker for Alzheimer’s disease and other brain disorders. By effectively managing variability in multisite, multi-scanner neuroimaging studies, our approach enhances the precision of atrophy measurements for clinical trials and longitudinal studies. This advancement promises to improve diagnostic and prognostic capabilities for neurodegenerative disorders.

## 1. Introduction

The quantification of brain atrophy through precise image registration is a critical component in the diagnosis and monitoring of neurodegenerative disorders such as Alzheimer’s disease, etc. [6]. However, the heterogeneity of neuroimaging data, stemming from variations in hardware, acquisition protocols, and imaging modalities, presents significant challenges to the development of robust and generalizable learning-based image registration techniques. These challenges are further compounded by the complex, non-linear deformations inherent in brain structures and the subtle nature of atrophic changes over time.

Image registration is an ill-posed problem and extensively studied. Traditional methods are iterative, e.g. Elastix [7], and ANTs [1] are some of the traditional registration tools. The methods provided in these tools are well-developed and provide both rigid and non-rigid registration capabilities. Classical iteration-based image registration methods, while effective in controlled settings, may falter when confronted with multi-site, multi-modal data. Pixel-based or intensity-based metrics, such as cross-correlation, and least squares, rely on the similarity of pixel intensities to align images accurately. However, this assumption may not always hold in real-world applications due to various factors such as differences in scanner characteristics, or temporal changes. Consequently, classical iterative methods as well as learning-based methods can be sensitive to outliers and intensity variations, which can affect the accuracy of the registration [10]. This limitation is somehow catered to by using mutual information or feature-based similarity metrics. This limitation of processing speed of these methods due to their iterative nature and ability to handle large variations in intensities or geometries has spurred interest in leveraging artificial intelligence, particularly deep learning approaches, to address these challenges. Recent advancements in AI-driven image registration have shown promise, yet the issue of domain invariance, the ability to perform consistently across different data sources and modalities, remains a critical area for improvement. Our work addresses this gap by introducing an AI-driven approach that is trained by using SSIM-based [11] loss function for a learning-based image registration model. This approach is specifically designed to handle the structural, intensity, and domain variances prevalent in 3D brain imaging data. By focusing on structural similarity rather than purely intensity-based metrics, our model achieves greater robustness to the variations inherent in multi-site neuroimaging studies.

These technical advancements collectively enable our model to achieve better accuracy in quantifying brain atrophy across a diverse range of 3D MRI data. The ability to accurately measure atrophy, a key biomarker for neurodegenerative diseases, has significant implications for both clinical practice and research. It allows for more precise tracking of disease progression, evaluation of treatment efficacy, and potentially earlier diagnosis of conditions like Alzheimer’s disease. In the following sections, we detail our methodology, present comprehensive experimental results, and discuss the implications of our work for the fields of medical image analysis and AI-driven healthcare diagnostics.

## 2. Methods

Our end-to-end image registration framework incorporates a multi-stage pipeline designed to address both global and local alignment challenges in 3D medical image analysis. Fig. 1 illustrates the overall block diagram of the registration process. The pipeline consists of three main stages:

**Figure 1.**
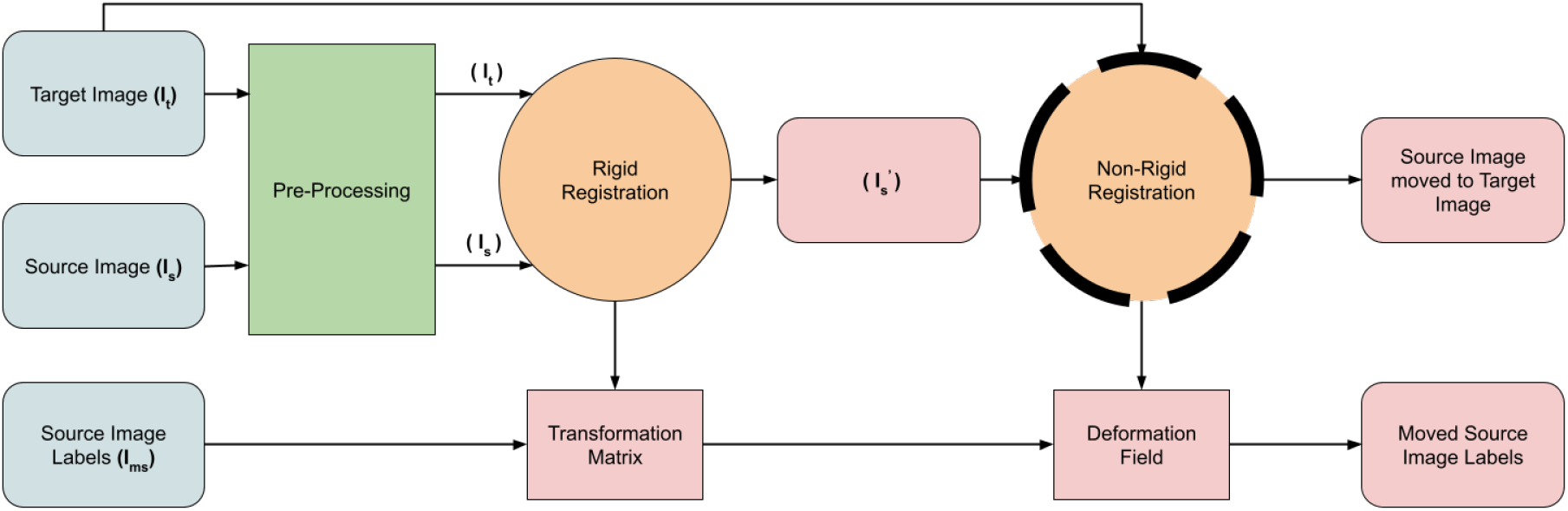
Standard pipeline for medical image registration: After pre-processing, images are affinely registered followed by non-rigid registration. Our work in this study focuses primarily on non-rigid registration

### 2.1. Pre-processing

Providers of the datasets have processed 3D data using N3 intensity normalization and standard bias field correction available in Freesurfer.

### 2.2. Global Alignment Module

This stage implements rigid registration to bring the source and target volumes into a common spatial reference frame. We employ ANTs [1] to first affinely align the pairs. The resultant transformation matrix is also applied to the labels. The transformation includes translation, rotation, scaling, and shearing:

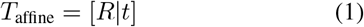

where *R* is a 3×3 rotation matrix and *t* is a 3×1 translation vector. The iterative method optimizes these parameters using a combination of mutual information loss and a regularization term:

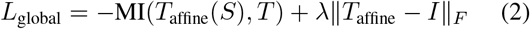

where MI is mutual information, *S* is the source volume, *T* is the target volume, *I* is the identity matrix, and *λ* is a regularization weight.

### 2.3. Non-rigid Registration Module

Following the global alignment, we apply our non-rigid registration network to address local deformations. This module utilizes the encoder-decoder architecture with a Spatial Transformer Network (STN) [5] as previously described. The key components are:

1. Feature Extraction: A 3D U-Net-like structure extracts multi-scale features from both globally aligned source and target volumes.
2. Deformation Field Estimation: The decoder generates a dense 3D displacement field *ϕ*(*x*) which is optimized over a loss function.
3. STN-based Warping: The displacement field is applied to the source volume via the STN.
4. Loss Computation: We use our custom 3D implementation of SSIM-based loss along with a regularization term.

The entire non-rigid pipeline is trained end-to-end. After getting rigid alignment, a non-rigid registration network is trained to optimize its parameters for the registration task. Our framework achieves state-of-the-art performance in handling multi-site 3D medical image registration data, particularly in the context of brain atrophy quantification for neurodegenerative disorders.

Our proposed architecture in Fig. 2, leverages an encoder-decoder framework to learn robust feature representations across source and target 3D volumetric data. The encoder comprises a series of 3D convolutional layers with residual connections, employing instance normalization and LeakyReLU activations to capture hierarchical features while maintaining spatial information. The decoder uses transposed 3D convolutions to upsample the encoded features, incorporating skip connections from the encoder to preserve fine-grained details.

**Figure 2.**
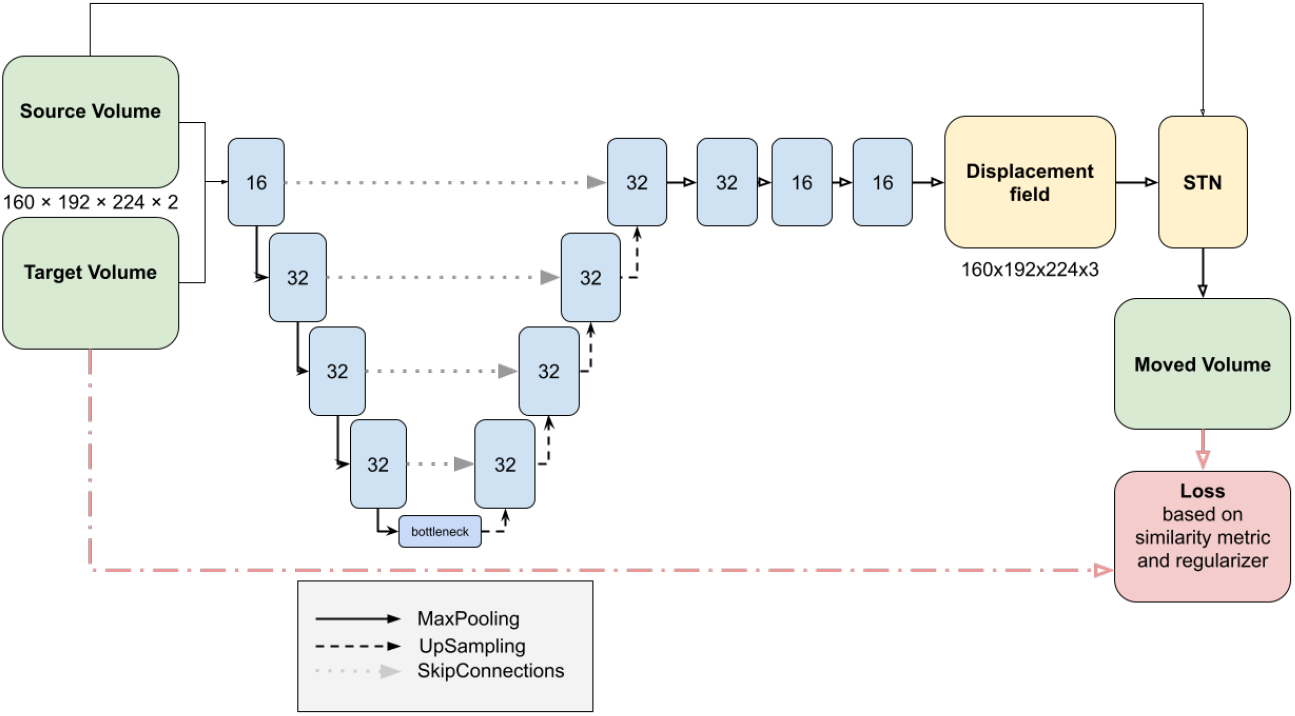
Overview of the deep learning-based image registration using a U-Net-based architecture as a backbone. The source image is transformed using the displacement field *ϕ* and the loss is computed between the target volume and the transformed source volume.

The output of the decoder is a dense 3D displacement field, *ϕ*(*x, y, z*), which represents the voxel-wise deformations required to align the source volume with the target. This displacement field is then fed into a Spatial Transformer Network (STN) module, specifically adapted for 3D volumetric data. STN applies the learned spatial transformations to the source volume, generating the Moved Volume *M* (*x*). Mathematically, this transformation can be expressed as :

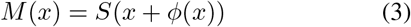

We used an SSIM-based loss function, with a window size of 11, to guide the learning process:

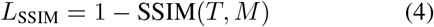

By choosing different hyperparameter values of the loss function, it can prioritize structural coherence over voxelwise intensity matching, making it more robust to interscanner and intra-modality variations in brain imaging datasets. Additionally, we incorporate a deformation regularization term to ensure smooth and plausible transformations:

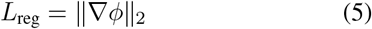

The total loss is a weighted combination:

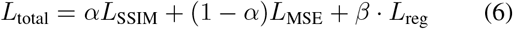

where *α* and *β* are empirically determined weighting factors. We use the Adam optimizer with a cyclical learning rate schedule to facilitate convergence and escape local minima in the high-dimensional parameter space of our 3D registration problem

Our work, extended from [9], focuses on applying robust 3D brain image registration techniques to map brain disorders. We use two large-scale 3D brain volume datasets i.e. OASIS [8] and ABIDE [3]. All brain volumes are resampled to 256×256×256 size and affine registration is performed to a common scan and cropped to the 160×192×224 size. We use 30 labeled structures as perused in [2] and we utilize 1854 scans from the OASIS dataset for training, 193 MRI scans from OASIS, and 103 MRI scans from ABIDE for evaluation purposes. Our model is trained using an SSIM-based loss function. As a simple example, Fig. 3 shows the displacement fields for a sample axial brain slice, we can see the reduced number of voxel folding for our SSIM-based loss function as compared to the MSE, given that our model is performing better than the baseline

**Figure 3.**
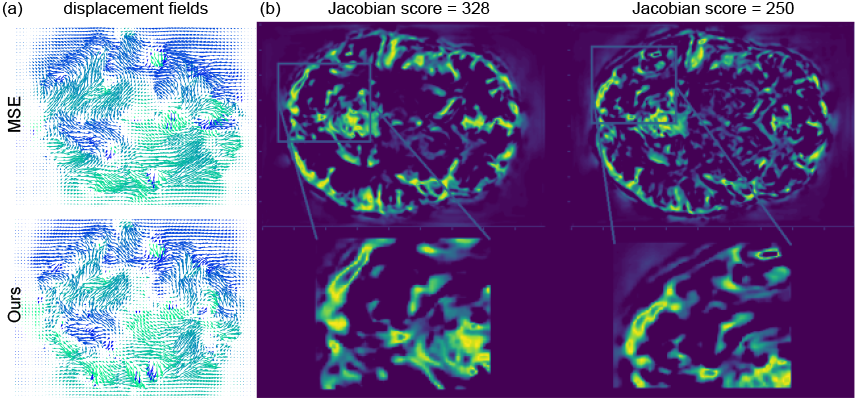
(a) Deformation fields for sample axial slice from 3D sample. (b) Jacobin score 250 with our (SSIM-based) loss function and 328 with the MSE-based loss

## 3. Results

In this section, we show the application of the proposed registration-based segmentation on the data from a new distribution i.e. ABIDE dataset. It can be noted that the prediction of our model is much closer to the ground truth labels as compared to the baseline (VoxelMorph) [2] model. Also, performance for dementia detection is presented. We first calculate the volumes of different brain regions and their variability across different cases from the ground-truth data. We then observe the same trend in the change of volumes using the proposed deep learning-based registration.

### 3.1. Performance on New Site Data

Fig. 4 presents the DICE scores for each structure as a box plot for completely new data from ABIDE [3]. For summarising purposes, we average the DICE scores of the left and right hemispheres. Our approach gets better results on almost all of the structures as compared to the baseline, significantly on ventral-dc, choroid plexus, lateral ventricle, and several other regions.

**Figure 4.**
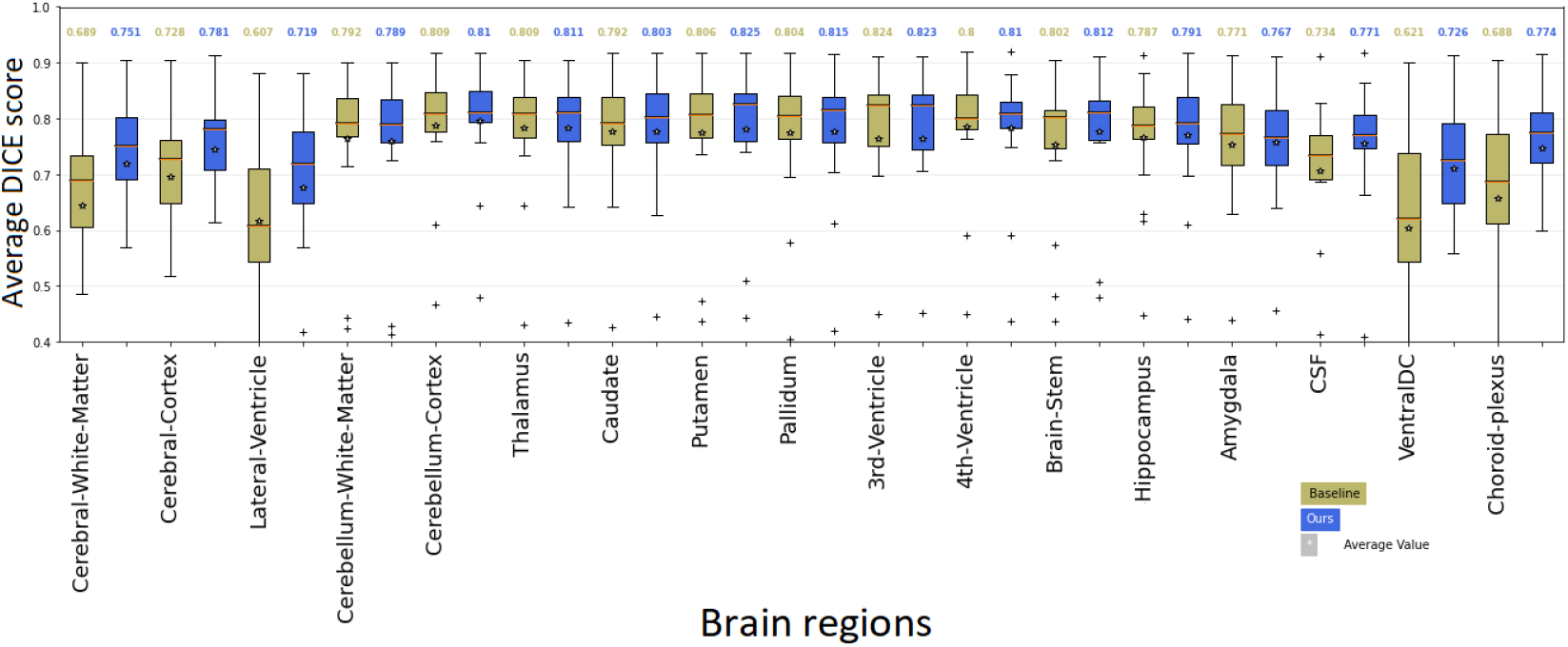
Boxplots of DICE scores for various anatomical structures for the baseline (VoxelMorph) and our model for unseen distribution data i.e. ABIDE. The left and right brain hemispheres are averaged together.

### 3.2. Data Preparation for Dementia Study

We use a publicly available OASIS [8] dataset for this purpose. Segmentation information is captured from the volumes preprocessed by the FreeSurfer [4]. A CSV file is provided for each subject’s visit with a clinical dementia rating (CDR). The actual usable MRI data for this study is less than the instances provided in the clinical spreadsheet, as dates for MRI scans and the dates for clinical visits/ratings are not aligned. For demented subjects, we take only those MRI scans that are scanned after the date when the clinical dementia rating *CDR >* 0 is given in the clinical record, and for control subjects, we take the MRI scans before the date when the clinical dementia rating *CDR* = 0 is given. Firstly, we get all the MRI scans that were never converted to dementia and make an average Atlas from these short-listed brains. All brains are affinely registered and then non-rigidly aligned and averaged. Subsequently, labels are computed for averaged Atlas by registering it to a normal healthy brain. In Table 1, the number of MRI scans is expressed for each level of CDR.

**Table 1.** Number of Sorted MRI Scans based on MRI Scan Time vs Clinical Diagnosis Time.

| Clinical Dementia Rating (CDR) | 0 | 0.5 | 1 | above 1 |
| --- | --- | --- | --- | --- |
| Number of MRI Scans | 728 | 442 | 84 | 6 |

### 3.3. CDR Rating and Change in Brain Regions’ Volume

Labels generated by FreeSurfer are used to calculate the volume for each region of control and demented subjects. Fig. 5 presents the average volume of each region against different CDR scores. It can be observed that except for a few, all other regions relatively remain unchanged.

**Figure 5.**
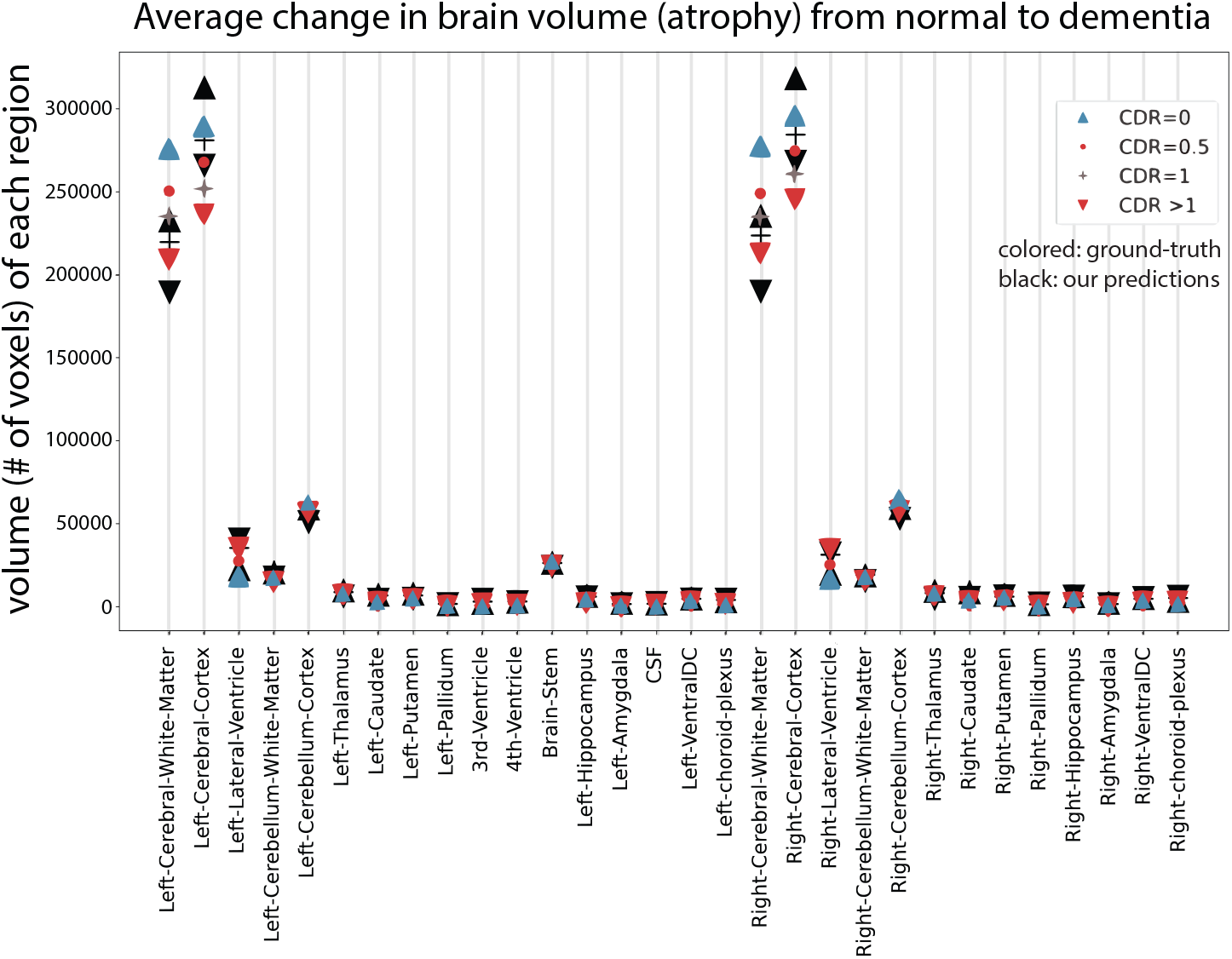
Average change in volume of different regions from Normal to Demented as per ground truth. Black shows our prediction and color shows the ground truth.

To test the effectiveness of both the baseline and our method for the prediction of brain regions, we apply registration models for atlas-based segmentation. Which, after finding the transformations from Atlas to Subject, the Atlas’ labels are also transformed to Subject space. Fig. 5 presents the prediction of volumes of different brain regions by our method in black markers. Our approach is providing the nearest results to the ground truth volumes. We observe that three regions from both the left and right hemispheres are affected by dementia. We calculate the difference between the predicted and ground-truth volumes of these three regions, as shown in Fig. 6. It can be observed that our predicted volumes are much closer to the ground-truth volumes compared to the predictions of the baseline method, whereas lower values (differences) are better. From this experiment, it is concluded that our method is able to quantify the atrophies in brain regions caused by neurodegenerative diseases with high precision. Our approach outperforms the baseline model in predicting the volumes of such brain regions, also demonstrating robustness for multisite data.

**Figure 6.**
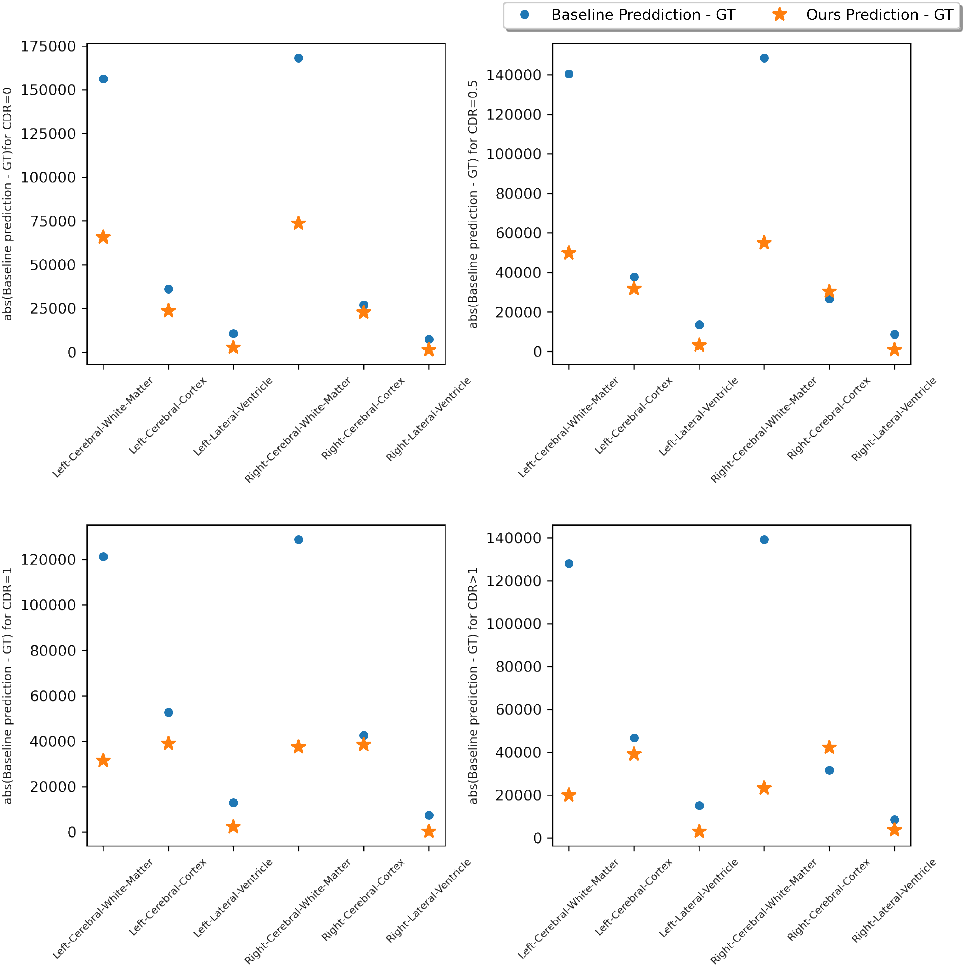
Average change in volume of different regions between predicted and ground-truth labels, difference of our prediction and ground-truth labels is less than the difference between the baseline and ground-truth labels

## 4. Conclusion

Our study demonstrates the effectiveness of a structure-based loss function in deep learning-based registration models for brain imaging. The results reveal that our method exhibits enhanced robustness when applied to novel, unseen multi-site data, a crucial advancement for clinical applications. We present a compelling case study on predicting volumetric changes in various brain regions affected by dementia, showcasing the potential of our approach in neurodegenerative disease research. These findings open promising avenues for future research, particularly in refining atlas-based segmentation processes. The implementation of multi-atlas label fusion (MALF) could further enhance the accuracy of neurodegenerative disease detection, potentially revolutionizing early diagnosis and treatment monitoring. By bridging the gap between advanced AI techniques and clinical neurology, our work contributes to the ongoing efforts to improve patient outcomes in the face of challenging neurological disorders.

